# Electrophysiological dependent antiarrhythmic drug response in population-based models of paroxysmal atrial fibrillation

**DOI:** 10.64898/2026.09.18.752633

**Authors:** Violeta Puche-García, Laura Martinez-Mateu, David Filgueiras-Rama, Lucía Romero, Javier Saiz

## Abstract

Response to antiarrhythmic drugs varies markedly across patients with atrial fibrillation (AF), suggesting that treatment efficacy depends on the interaction between drug-specific mechanisms and patient-specific electrophysiological substrate. Here, we used population-based computational models to investigate how electrophysiological substrate and inter-individual ionic variability influence pharmacological efficacy and the underlying mechanisms.

Two populations of human atrial models were generated from distinct substrates: a reference left atrial model and a second model incorporating inward-rectifier-enhancement (IRE) through a 2-fold increase in I_K1_ and I_K,ACh_. Both populations were independently calibrated against the same experimental datasets from patients with paroxysmal AF (pAF), yielding pAF and IRE-pAF populations. Sustained reentrant activity was induced in two-dimensional tissue simulations and subsequently used to assess cardioversion efficacy of flecainide, vernakalant and tertiapin-Q.

Despite satisfying the same calibration criteria, IRE-pAF population exhibited a more arrhythmogenic phenotype: shorter refractoriness, higher dominant frequency (DF) and greater rotor stability. Antiarrhythmic efficacy markedly differed between substrates. Flecainide cardioversion decreased in IRE-pAF compared with pAF (34% vs 21%), whereas I_K,ACh_-targeting therapies preserved or improved efficacy in IRE-pAF (vernakalant: 41% vs 44%, tertiapin-Q: 11% vs 18%). Across drugs and substrates, rotor DF strongly influenced cardioversion outcome, with higher-frequency rotors showing lower termination rates. Drug-induced DF reduction emerged as a key mechanism associated with successful cardioversion, whereas effective refractory period (ERP) prolongation alone did not consistently explain treatment efficacy. In pAF, vernakalant achieved higher cardioversion efficacy than flecainide despite a smaller increase in ERP and greater DF reduction. Ionic analyses further showed that elevated I_K,ACh_ favored responses to vernakalant and tertiapin-Q.

These findings demonstrate that cardioversion efficacy emerges from the interaction between electrophysiological substrate, rotor dynamics and drug-specific mechanisms. In particular, substrates differing in inward rectifier activity exhibit distinct response patterns, while DF emerges as a robust marker of pharmacological susceptibility and a potential guide for drug-mediated AF termination.

**Author summary:** Antiarrhythmic drugs are commonly used to restore normal rhythm in patients with paroxysmal atrial fibrillation (pAF), but their effectiveness varies substantially among individuals. The reasons underlying this variability remain incompletely understood, limiting our ability to select the most appropriate treatment for a given patient.

In this study, we used populations of computational models of human atrial tissue to investigate how differences in the underlying electrophysiological substrate influence drug response and the underlying mechanisms. We generated two pAF-like populations differing in the level of inward rectifier activity. Then, we evaluated the effects of three antiarrhythmic drugs with distinct mechanisms of action.

Our results show that, while some therapies became less effective in the inward-rectifier-enhanced substrate, others maintained or even improved their performance. We also found that rotor dominant frequency (DF) was closely linked to treatment success and that drugs producing greater DF reduction were more likely to terminate AF.

These findings highlight the importance of considering substrate-specific electrophysiology when evaluating antiarrhythmic therapies and illustrate how computational modeling can help uncover mechanisms of drug response. More broadly, our results suggest that electrophysiological markers such as DF may help identify patients more likely to benefit from specific rhythm-control strategies, supporting future personalized approaches for AF treatment.

## Introduction

Antiarrhythmic drugs remain a first-line rhythm control strategy for recent-onset and paroxysmal atrial fibrillation (pAF) [1, 2]. However, pharmacological response remains difficult to predict and varies substantially across AF episodes and patients, highlighting both our incomplete understanding of the mechanisms governing the arrhythmia and the challenge of identifying patient-specific determinants of treatment success [3]. Increasing evidence suggests that differential antiarrhythmic response emerges from the interplay between drug-specific actions [4], patient variability [3, 5] and the underlying electrophysiological substrate sustaining AF [6, 7]. Recurrent AF episodes progressively reshape atrial electrophysiology (”AF begets AF” [8]), promoting faster activation rates and more stable AF-maintaining mechanisms [9–11] that may fundamentally alter pharmacological response [12]. These changes arise partially from complex alterations in ionic current expression and regulation that contribute to the maintenance and progression of AF [6]. In particular, inward rectifier remodeling has attracted considerable interest due to its potential role in stabilizing high-frequency reentrant activity. Increased I_K1_ and constitutively active I_K,ACh_ have been associated with action potential shortening, rotor acceleration, increased dominant frequency (DF) and enhanced AF maintenance [10, 11, 13–15].

Atrial activation frequency has emerged as an important marker of AF organization and therapeutic response. Clinical and experimental studies have demonstrated that higher DF is associated with increased AF complexity and reduced cardioversion efficacy [11, 16–18]. Importantly, conventional antiarrhythmic drugs, including fast sodium current (I_Na_) and rapid delayed-rectifier potassium current (I_Kr_) blockers such as flecainide and ibutilide, preferentially terminate AF episodes with relatively low activation frequencies (*≈* 5.6 Hz), whereas their efficacy decreases in higher-DF episodes [19, 20]. These findings identify AF substrate complexity and activation rate as key modulators of pharmacological cardioversion efficacy.

The development of novel atrial-selective antiarrhythmic strategies has emerged as an attractive approach to improve AF termination [21]. Vernakalant combines preferential atrial I_Na_ blockade with additional effects on atrial-specific potassium currents, including I_K,ACh_, at therapeutic concentrations [22, 23]. Specifically, selective I_K,ACh_ inhibition has been proposed as a promising strategy to suppress high-frequency AF by destabilizing rapid atrial activation patterns [21]. Experimental studies using tertiapin-Q, a highly selective I_K,ACh_ blocker, demonstrated prolongation of atrial refractoriness and reduced AF maintenance [24], directly supporting selective I_K,ACh_ targeting as a potential antiarrhythmic strategy [25].

Nevertheless, despite evidence linking activation frequency and inward rectifier remodeling to antiarrhythmic response, how substrate-specific electrophysiological remodeling differentially modulates the efficacy of conventional and atrial-selective antiarrhythmic strategies remains unclear. In particular, the mechanisms underlying the differential response of high-frequency substrates to distinct pharmacological interventions are still poorly understood.

Computational modeling provides a powerful framework to address these questions because it enables systematic control of tissue electrophysiology and pharmacological interventions under reproducible conditions that are difficult to isolate experimentally or clinically [26]. Additionally, population-based methodologies allow the evaluation of inter-subject variability and heterogeneous drug responses across a broad range of electrophysiological conditions [5, 27, 28]. Such approaches are particularly suitable for investigating how inward rectifier remodeling and rotor DF alter antiarrhythmic efficacy and reshapes the mechanisms leading to successful cardioversion.

In the present study, we hypothesized that antiarrhythmic efficacy depends on the interaction between the underlying tissue electrophysiological properties during AF and drug-specific electrophysiological actions, resulting in patient-specific pharmacological responses. To test this hypothesis, we constructed two populations of human atrial action potential models reproducing pAF electrophysiology: one based on a reference left atrial model and the other incorporating enhanced inward rectifier remodeling through a 2-fold increase in I_K1_ and I_K,ACh_. Using tissue-scale simulations derived from these populations, we evaluated substrate-dependent responses to three compounds (flecainide, vernakalant, tertiapin-Q) and investigated the mechanisms underlying successful cardioversion. Our findings provide mechanistic insights into how AF substrate properties and drug-specific effects interact to determine antiarrhythmic efficacy, supporting substrate-dependent approaches for AF pharmacological cardioversion.

## Methods

### Populations of human paroxysmal atrial models

The Courtemanche–Ramirez–Nattel model [29] was selected to reproduce the human atrial action potential since it has been extensively used to model and study the action of antiarrhythmic drugs [5, 27, 28, 30, 31].

The model was extended to incorporate additional ionic currents to improve representation of atrial-specific electrophysiology. In particular, the acetylcholine-activated potassium current (I_K,ACh_) was incorporated following Grandi et al. formulation [32] using a basal acetylcholine concentration of 0.005 *µ*M [33, 34] to reproduce cholinergic modulation of atrial repolarization [10, 35] and enable simulation of pharmacological I_K,ACh_-targeting (vernakalant and tertiapin-Q). Additional formulations included the small-conductance Ca^2+^-activated potassium current 67 (I_SK_) [36], the two-pore domain potassium current (I_K2P_) [37] and stretch-activated channels (I_SAC_) [38]. Model re-parameterization was performed to preserve baseline action potential duration and morphology after incorporation of the additional ionic currents. Details of the incorporation of the additional ionic currents, subsequent model re-parameterization and validation procedures are provided in the Supplementary Methods (S1 Appendix).

To reproduce left atrial electrophysiology, I_Kr_ was increased by 60% [5, 34, 39], accounting for the reported reduction in action potential duration (APD) between the right and left atria [40]. This left-atrial formulation was used as the reference model. From this, an additional two-fold increase in I_K1_ and I_K,ACh_ was introduced to generate an inward-rectifier-enhanced (IRE) model. These currents were selected based on experimental and clinical evidence linking inward-rectifier remodeling to maintenance and progression of pAF. Increased I_K1_ has been associated with action potential shortening, rotor acceleration and DF increase during AF progression [14, 15]. Likewise, enhanced I_K,ACh_ activity has been related to higher rotor DF and left atrial DF gradients, particularly in paroxysmal AF substrates [13, 15]. Therefore, simultaneous enhancement of both inward-rectifier currents was used to reproduce a more arrhythmogenic pAF substrate characterized by faster and more stable reentrant activity. Consequently, two distinct baseline conditions were defined: a left atrium pAF substrate (pAF) and a left atrium inward-rectifier-enhanced pAF substrate (IRE-pAF).

To account for electrophysiological variability, populations of models were generated as described by Sánchez de la Nava et al. [27]. A total of 10,000 ionic profiles were created using Latin Hypercube Sampling by varying 13 selected ionic conductances from -50% to +100%, including the ultrarapid, rapid and slow delayed rectifier K^+^ current (G_Kur_, G_Kr_ and G_Ks_, respectively), transient outward K^+^ current (G_to_), inward rectifier K^+^ current (G_K1_), L-type Ca^2+^ current (G_CaL_), fast Na^+^ current (G_Na_), Na^+^+/K^+^ pump (G_NaK_), Ca^2+^/Na^+^ exchanger (G_NCX_) and the additional ionic currents G_K,ACh_, G_SK_, G_K2P_ and G_SAC_. The same set of 10,000 ionic scaling combinations was applied to both baseline substrates to generate the two populations.

Population calibration was performed at both cellular (0D) and tissue (2D) levels and the acceptance ranges are summarized in Table 1. First, all ionic profiles were paced for 50 beats at single-cell level [5, 41] to evaluate action potential (AP) biomarkers at 1 Hz [42, 43] and calcium transient biomarkers at 2 Hz [44]. AP calibration was based on resting membrane potential (RMP) and APD at 90%, 50% and 20% repolarization (APD_90_, APD_50_, and APD_20_) data obtained from recordings of the human left atrial appendage cardiomyocytes from patients with paroxysmal AF reported by Verkerk et al. [42] and Casini & Verkerk et al. [43]. Calcium transient calibration was based on the relative variability of diastolic (CaT_diast_), systolic (CaT_sys_) and amplitude (CaT_amp_) calcium concentration data, together with the calcium transient decay time constant (*τ_decay_*), obtained from human right atrial appendage cardiomyocytes from patients with paroxysmal AF reported by Voigt et al. [44]. Experimental coefficients of variation were applied to baseline simulated values to define physiologically constrained acceptance ranges. AP and calcium transient acceptance ranges were generally defined as mean *±* 2 standard deviations (SD) from the corresponding experimental datasets. When multiple studies were available for the same biomarker, the widest interval was retained to account for inter-study variability. Models satisfying the cellular electrophysiological criteria were subsequently simulated in 2D tissue to evaluate rotor DF under arrhythmic conditions (see Tissue simulations section). Based on clinical data reported by Atienza et al. (5.7 *±* 0.8 Hz) [11], the acceptable DF values ranged from 4.1 to 7.3 Hz (mean *±* 2 SD). Profiles exhibiting DF values outside this range were discarded. Additional details regarding calibration procedures and acceptance ranges are provided in the Supplementary Methods (S1 Appendix).

**Table 1.** Biomarker acceptance ranges used for calibration of both atrial populations. Experimental and clinical biomarker acceptance ranges used for calibration across both populations at the cellular (0D) and tissue (2D) levels. Abbreviations: APD: Action Potential Duration; RMP: Resting Membrane Potential; DF: Dominant frequency.

| Biomarker | Min Value | Max Value | Simulation | Reference |
| --- | --- | --- | --- | --- |
| APD<sub>90</sub> (ms) | 160 | 254 | 0D-level 50 pulses at 1Hz | Verkerk et al. [42], |
| APD<sub>50</sub> (ms) | 0 | 48 |  | Casini et al. [43] |
| APD<sub>20</sub> (ms) | 1 | 5 |  |  |
| RMP (mV) | -93 | -70 |  |  |
| CaT<sub>diast</sub> (nM) | 81 | 172 | 0D-level 50 pulses at 2Hz | Voigt et al. [44] |
| CaT<sub>sys</sub> (nM) | 394 | 622 |  |  |
| CaT<sub>amp</sub> (nM) | 275 | 488 |  |  |
| $\tau_{decay}$ (ms) | 119 | 176 | | |
| DF (Hz) | 4.1 | 7.3 | 2D-level S1-S2 protocol | Atienza et al. [11] |

Only models satisfying all calibration criteria were retained, resulting in 853 models in the pAF population and 282 models in the IRE-pAF population. The complete sets of ionic scaling factors defining the calibrated populations are provided in the Supporting Information, including all profiles from the pAF population (S1 Data) and all profiles from the IRE-pAF population (S2 Data).

### Tissue simulations: vulnerability, rotor induction and reentrant dynamics

Prior to tissue simulations, each profile in both populations was paced for 50 beats at a cycle length of 1000 ms in a single-cell environment [5, 41].

Fiber simulations were carried out to characterize tissue electrophysiology. Effective refractory period (ERP) and conduction velocity (CV) were computed for all calibrated profiles using a 5-cm-long narrow tissue domain, discretized with the same spatial resolution (300 *µ*m) and electrophysiological properties used in the subsequent 2D simulations. The fiber consisted of two elements across its width, resulting in a tissue domain of 600 *µ*m × 5 cm. ERP and CV were determined using stimulation protocols adapted from those previously described by Dasi et al. [5]. Wavelength (WL), an index of reentrant vulnerability representing the distance traveled by the activation wave during refractoriness, was calculated as the product of ERP and CV (WL = ERP × CV).

Two-dimensional tissue simulations were performed to characterize vulnerability and assess pharmacological cardioversion. As in previous studies [14, 33, 45, 46], this simplified tissue model enables isolation of ionic mechanisms from additional structural and electrophysiological heterogeneities. Simulations were carried out on a 5 × 5 cm^2^ atrial patch discretized with a spatial resolution of 300 *µ*m. Tissue conductivity parameters were adjusted to reproduce physiological atrial conduction velocities (60–90 cm/s), using a longitudinal conductivity coefficient of 0.19 S/m and an anisotropy ratio of 0.35 [33, 47]. Electrical propagation was simulated using the monodomain equation with the Elvira software [48].

Reentrant activity was induced using an S1–S2 cross-field stimulation protocol [14, 33, 45, 46]. After an initial stabilization phase consisting of 10 planar beats (S1) at a basic cycle length of 1000 ms, a rectangular premature stimulus (S2) was applied in the lower-left corner using different coupling intervals to induce reentry during 6-seconds-long simulations. Vulnerability was quantified as the width of the vulnerable window, defined as the range of S2 coupling intervals inducing sustained reentry, using a temporal resolution of 1 ms. Rotor inducibility was defined as the initiation of at least one complete reentrant rotation, whereas sustained activity was defined as reentry lasting longer than 5 seconds [33]. When multiple S2 coupling intervals within the vulnerable window induced sustained reentry (i.e., in profiles with wider vulnerable windows), representative rotors were selected to avoid over-representation of these profiles. Specifically, three rotors were selected for vulnerable windows *≥* 3 ms (beginning, middle and end of the vulnerable window), two for windows of 2 ms and one for vulnerable windows of 1 ms. Only selected sustained rotors were considered for subsequent pharmacological and mechanistic analyses.

Rotor dynamics were analyzed using phase mapping based on the Hilbert transform of transmembrane voltage signals [33, 47]. Rotor tip trajectories were computed by detecting phase singularities and meandering area was estimated using an ellipse approximation. Rotor DF was computed from nine pseudo-electrogram recording locations distributed across the tissue and the median value was used as the DF measurement for each rotor simulation. For profiles with multiple representative rotors selected from different S2 coupling intervals within the vulnerable window, DF and meandering area were quantified as the median across representative rotors, as similar rotor dynamics were observed regardless of S2 coupling interval.

Rotor DF was further used to stratify ionic profiles into different dynamical regimes. While population calibration was based on the mean *±* 2 SD of the clinical DF distribution reported by Atienza et al. [11], the mean *±* SD thresholds were used to define low- and high-DF subgroups (4.1–4.9 Hz and 6.5–7.3 Hz, respectively).

### Pharmacological modeling and cardioversion protocol

Sustained rotors were subjected to pharmacological interventions to evaluate cardioversion success. Drug effects were applied after 5 seconds of sustained reentrant activity and simulations were continued for an additional 10 seconds. Cardioversion success was defined as termination of reentrant activity after drug application during the 10-second-lasting simulation. Cardioversion efficacy was computed as the proportion of terminated rotors relative to the total number of sustained rotors.

Three compounds were evaluated: flecainide, a class Ic antiarrhythmic drug; vernakalant, an atrial-selective antiarrhythmic compound; and tertiapin-Q, a selective I_K,ACh_ blocker. Drug effects were modeled using a simple pore-block formulation determined by IC_50_ values and Hill coefficients [5, 27, 28]:

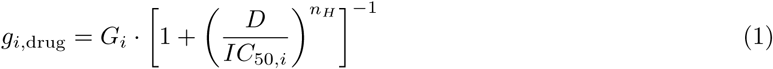

where, for current *i*, *G_i_* is the maximum conductance, *IC*_50*,i*_ is the half-maximal inhibitory concentration, *n_H_* is the Hill coefficient and *D* is the free drug concentration.

Because of the variability among published IC_50_ values, a preliminary calibration of flecainide and vernakalant formulations was performed to reproduce experimentally reported electrophysiological effects. Following previous methodologies [49–51], candidate IC_50_ values for each affected current were first collected from the literature and used to construct initial drug formulations. These formulations were subsequently evaluated through fiber simulations reproducing the cycle lengths and drug concentrations reported in the corresponding experimental studies. Simulated drug-induced changes in APD_90_ and maximum depolarization velocity were compared to published experimental observations and different combinations of reported IC_50_ values were explored. Final drug formulations were selected based on their overall agreement with the available electrophysiological responses reported experimentally. Additional details regarding the final IC_50_ values and comparison between simulated and experimental drug-induced electrophysiological effects are provided in the Supplementary Methods (S1 Appendix).

Tertiapin-Q was modeled as a selective I_K,ACh_ blocker using the experimentally reported IC_50_ value of 0.07 *µ*M and concentrations previously evaluated by Takemoto et al. [52]. No additional calibration procedure was performed.

Two concentrations within the reported therapeutic plasma concentration range were evaluated for each compound: flecainide (1.5 and 3 *µ*M) [53], vernakalant (10 and 30 *µ*M) [54] and tertiapin-Q (0.1 and 0.2 *µ*M) [52]. Resulting ionic current block percentages are summarized in Table 2.

**Table 2.** Ionic current blockade for each drug and concentration.

| Drug | Concentration | Ionic current blockade (%) |  |  |  |  |  |  |
| --- | --- | --- | --- | --- | --- | --- | --- | --- |
| | | $I_{CaL}$ | $I_{Kr}$ | $I_{Ks}$ | $I_{Kur}$ | $I_{to}$ | $I_{Na}$ | $I_{K,ACh}$ |
| Flecainide | 1.5 $\mu M$ | 6 | 50 | 7 | 34 | 22 | 25 | |
| | 3 $\mu M$ | 11 | 65 | 13 | 51 | 31 | 41 | |
| Vernakalant | 10 $\mu M$ | 19 | 33 | | 40 | 40 | 17 | 50 |
| | 30 $\mu M$ | 42 | 60 | | 67 | 67 | 38 | 75 |
| Tertiapin-Q | 0.1 $\mu M$ | | | | | | | 59 |
| | 0.2 $\mu M$ | | | | | | | 74 |

For each calibrated profile, ERP, CV and WL were also computed under each treatment using fiber simulations (see Tissue simulations section). Biomarkers related to rotor dynamics (DF and meandering area) were also computed after drug administration, considering only non-terminated rotors. Drug-induced changes in fiber and rotor dynamics biomarkers were quantified as percentage differences relative to control conditions before drug application. For profiles exhibiting multiple rotors, representative values were summarized as the median per profile.

### Ionic analysis of pharmacological response

To identify ionic determinants of pharmacological response, distributions of ionic conductances were compared between responder and non-responder profiles for each drug and population. Responders were defined as profiles in which drug administration resulted in termination of at least one sustained rotor, whereas non-responders were defined as profiles in which reentry persisted after treatment. Conductance values were expressed as percentage variation relative to the corresponding baseline conductance used for population generation.

Additional analyses were performed on profiles exhibiting exclusive response to a single drug. Exclusive responders were defined as ionic profiles successfully cardioverted by only one compound among the three evaluated drugs. Distributions of selected ionic conductances were compared across exclusive responder groups to identify ionic signatures associated with differential drug efficacy.

## Results

### Inward-rectifier enhancement shifts calibrated pAF-like populations toward a more arrhythmogenic regime

Two populations of calibrated left-atrial pAF-like models were generated from two distinct baseline substrates. Although both were calibrated to the same experimental data (Table 1), pAF and IRE-pAF populations exhibited distinct ionic, cellular and tissue-level properties (Fig 1). From the 10,000 candidate sampled ionic combinations, 853 models fulfilled the calibration criteria in the pAF population and 282 in the IRE-pAF population. Representative action potential morphologies from the candidate and calibrated populations are shown in Fig 1A.

**Fig 1.**
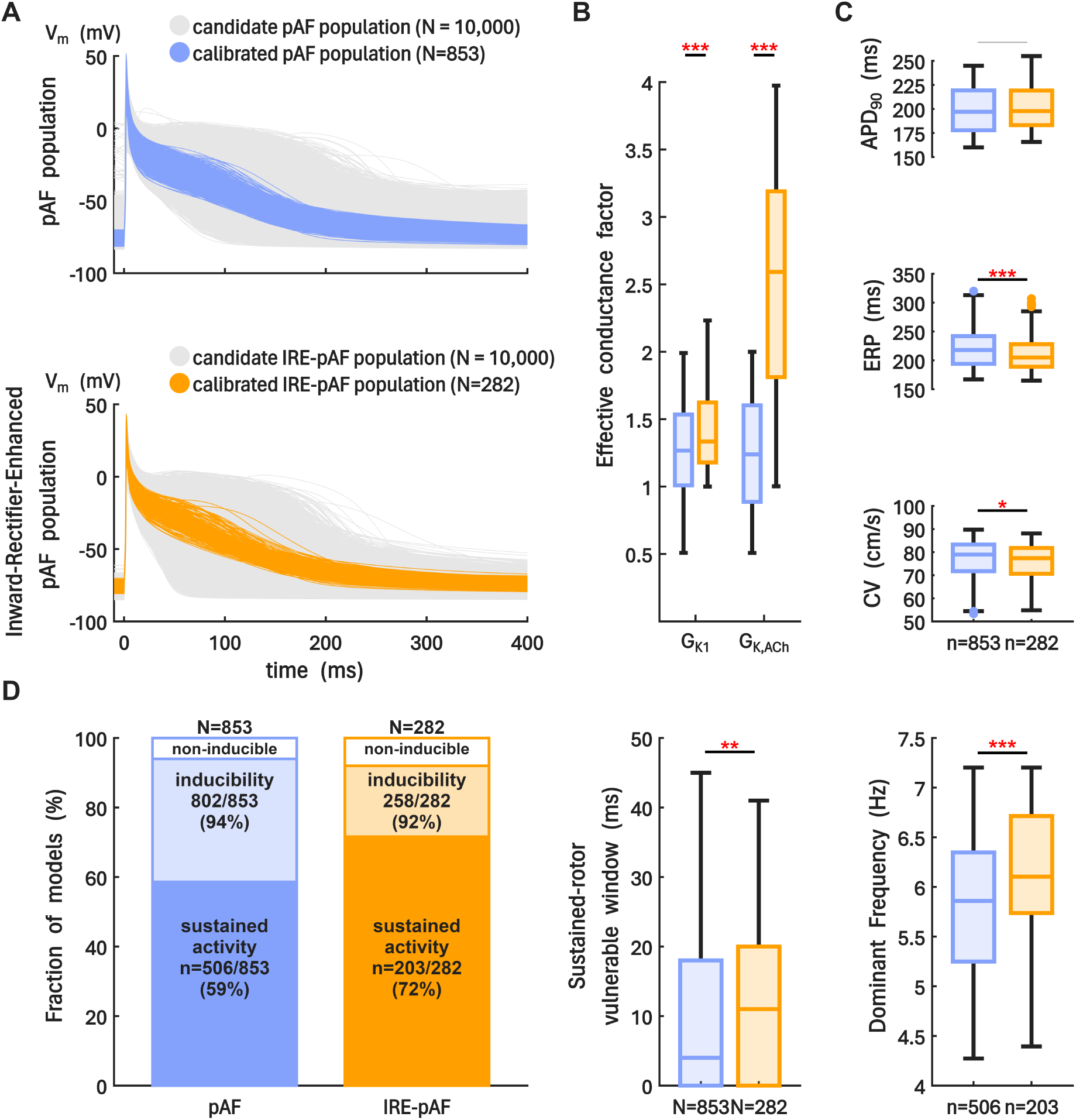
Electrophysiological and arrhythmogenic characterization of two calibrated pAF-like model populations. A: Action potential morphologies for both calibrated populations of models (pAF and IRE-pAF populations are shown in orange and blue, respectively). Both populations were independently calibrated to the same experimental data from paroxysmal AF. B: Ionic characterization based on the effective conductance distributions of the inward-rectifier currents I_K1_ and I_K,ACh_ after population calibration. C: Electrophysiological characterization at the cellular and tissue level, including APD_90_, effective refractory period (ERP) and conduction velocity (CV). D: Reentrant substrate characterization, including rotor inducibility, sustained activity (life span *>* 5 seconds), vulnerable window and dominant frequency (DF). Statistical comparisons were performed using the Wilcoxon rank-sum test: \**p <* 0.05, \*\**p <* 0.01, \*\*\**p <* 0.001.

Fig 1B summarizes the effective conductance distributions of the two inward-rectifier currents defining the electrophysiological substrate. Despite the two-fold increase in baseline I_K1_ and I_K,ACh_ used to construct the IRE-pAF substrate, calibration constrained the final conductance distributions and resulted in different effective shifts for both currents. Effective G_K1_ values exhibited a moderate displacement toward higher values in the IRE-pAF population, with substantial overlap between populations and similar upper ranges. In contrast, G_K,ACh_ showed a marked separation between populations, with an approximately two-fold increase in median values, displaced interquartile ranges and substantially higher maximum conductances in IRE-pAF. Thus, whereas calibration partially compensated for the imposed increase in G_K1_, the IRE-pAF population remained characterized by a substantially greater contribution of I_K,ACh_. Consequently, differences between populations after calibration were more pronounced for I_K,ACh_ than for I_K1_. The remaining ionic conductances exhibited broadly comparable ranges in both populations, indicating that calibration preserved substantial electrophysiological variability beyond the inward-rectifier currents. A detailed characterization of ionic variability across populations is provided in S1 Figure.

At the cellular and tissue level (Fig 1C), APD_90_ was comparable between the two populations (197 *±* 24 ms in pAF vs 198 *±* 23 ms in IRE-pAF, n.s.). Despite the similar APD_90_, the IRE-pAF population exhibited lower ERP and slightly slower CV than the pAF population (ERP: 205 *±* 29 vs 218 *±* 32 ms, *p <* 0.001; CV: 77 *±* 8 vs 79 *±* 8 cm/s, *p <* 0.05). Thus, inward-rectifier enhancement altered tissue refractoriness and conduction without producing a significant shift in APD_90_, suggesting a modest shift toward a more remodeled electrophysiological phenotype in the IRE-pAF population.

The impact of these cellular and tissue-level differences on reentrant behavior was subsequently assessed using 2D simulations and an S1–S2 rotor induction protocol (Fig 1D). The probability of inducing reentrant activity (at least one rotation) was similar for both populations (94% in pAF vs 91% in IRE-pAF). However, sustained rotor activity (rotor life span *>* 5 s) was more frequent in IRE-pAF than in pAF (72% vs 59% of calibrated models). Likewise, the vulnerable window for sustained activity was wider in IRE-pAF (11 *±* 11 ms) than in pAF (4 *±* 11 ms, *p <* 0.01), indicating a substrate more prone to sustaining long-lasting reentries. Rotor DF, although constrained to clinical observation, was also displaced to higher values in IRE-pAF compared to pAF (6.1 *±* 0.68 vs 5.9 *±* 0.72 Hz, *p <* 0.001). Altogether, these results indicate that inward-rectifier enhancement does not merely shorten repolarization, but also produces a faster, more vulnerable and more stable reentrant regime, consistent with a more arrhythmogenic substrate.

### Pharmacological cardioversion efficacy diverges across electrophysiological substrates

Pharmacological cardioversion (Fig 2) was evaluated exclusively in simulations exhibiting long-lasting reentrant activity. A total of 1366 sustained rotor episodes were analyzed in the pAF population and 564 in the IRE-pAF population, allowing direct assessment of the probability that an established reentrant episode terminates under pharmacological intervention.

**Fig 2.**
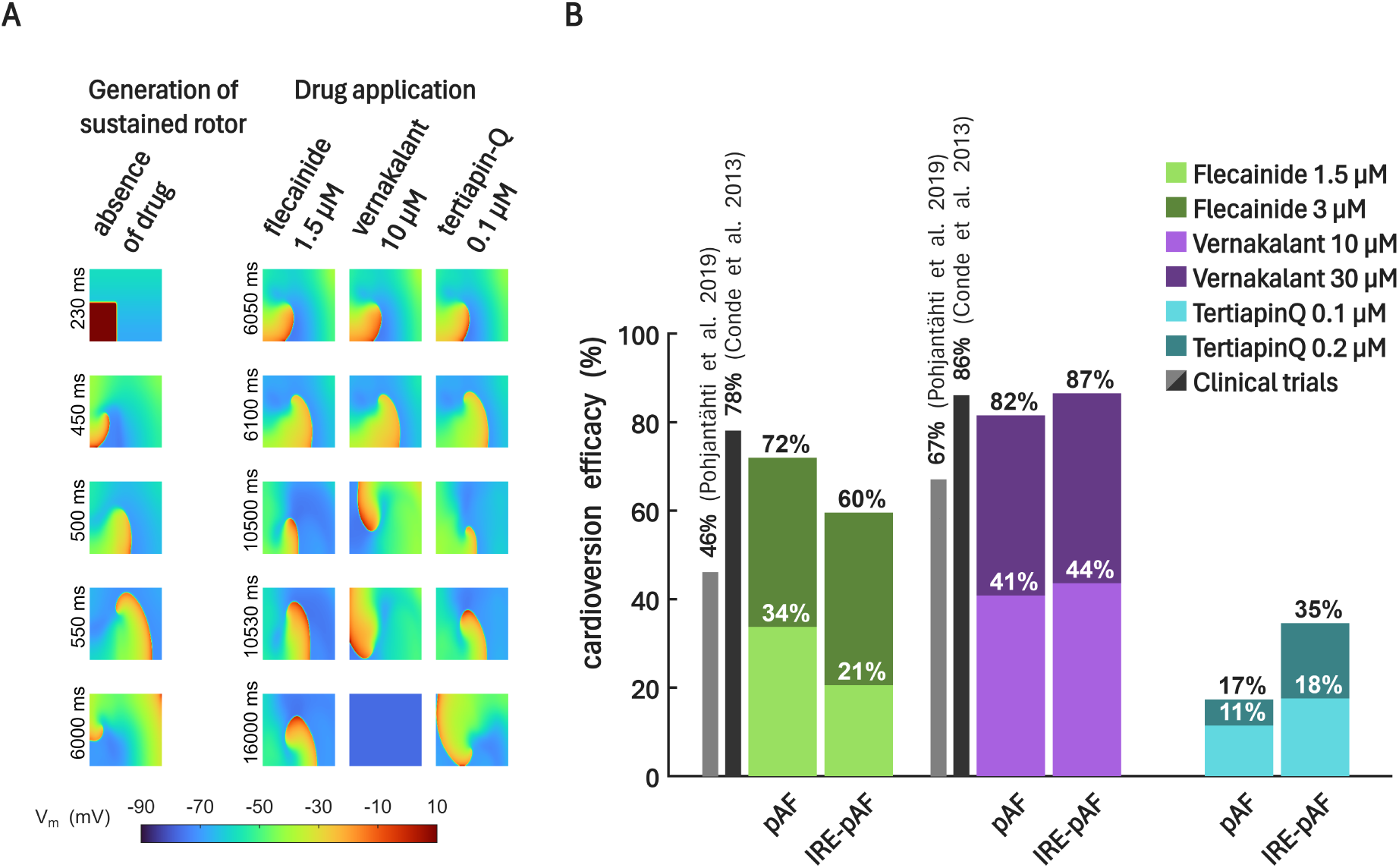
Pharmacological cardioversion study. A: Representative simulation showing rotor induction using an S1–S2 protocol (left column) and subsequent pharmacological intervention (right columns) for flecainide 1.5 *µ*M, vernakalant 10 *µ*M and 10 tertiapin-Q 10 *µ*M. B: Cardioversion efficacy (%) across sustained rotors in the two pAF-like populations. Cardioversion was defined as termination of sustained rotor activity during the simulation following drug administration and computed as the ratio of terminated rotors to total sustained rotors. In-silico results (color bars) are shown for low and high concentrations (lighter and darker shades, respectively) and are compared with reported human clinical values (black/gray bars).

For a specific ionic profile, Fig 2A illustrates a representative simulation of rotor induction and subsequent pharmacological interventions, highlighting differences in termination between drugs (in this case, vernakalant resulted in rotor termination, whereas reentrant activity persisted under flecainide and tertiapin-Q).

Analysis of all sustained rotors (Fig 2B) revealed that pharmacological cardioversion differed between drugs and depended on the underlying electrophysiological substrate. Across populations and concentrations, vernakalant consistently achieved the highest cardioversion rates, followed by flecainide and tertiapin-Q, reproducing the efficacy hierarchy reported in clinical studies. However, the electrophysiological substrate affected each drug differently. Flecainide lost cardioversion success in IRE-pAF relative to pAF at both concentrations (by 13 and 12 percentage points at low and high concentration, respectively), whereas vernakalant slightly improved its performance and tertiapin-Q exhibited a clearer gain in the IRE-pAF substrate. This divergence became particularly evident at high concentration. In pAF, the difference between flecainide and vernakalant was 10 percentage points (72% vs 82%), whereas in IRE-pAF it increased to 27 percentage points (60% vs 87%). The largest electrophysiological substrate-dependent increase was observed for tertiapin-Q, whose antiarrhythmic performance rose from 17% in pAF to 35% in IRE-pAF.

As expected, termination rates increased with concentration for all drugs and in both populations. However, the magnitude of this gain differed across compounds. Flecainide and vernakalant each showed an increase of approximately 40 percentage points from low to high concentration in both substrates, whereas tertiapin-Q showed a smaller increase in pAF and a more moderate increase in IRE-pAF. Thus, the greater dose-dependent improvement observed for flecainide and vernakalant may be associated with their broader multi-channel blockade compared to the selective I_K,ACh_ inhibition of tertiapin-Q. Particularly, despite similar levels of I_K,ACh_ blockade at high concentration (75% for vernakalant vs 74% for tertiapin-Q), vernakalant achieved substantially higher cardioversion efficacy, suggesting that combined modulation of I_K,ACh_ and additional ionic currents provides a greater antiarrhythmic effect than selective I_K,ACh_ blockade alone.

### Rotor deceleration explains cardioversion efficacy across drugs and electrophysiological substrates

To investigate the electrophysiological mechanisms underlying these efficacy differences, we quantified drug-induced changes in DF and ERP as shown in Fig 3 for low concentrations (see S2 Figure for high concentration drug-induced changes). Broadly, the three drugs induced both DF reduction and ERP prolongation (Fig 3B and Fig 3C, respectively). Fig 3A, left panel, displays a representative example of changes in both the rotor meandering area and DF for a given sustained rotor before (first row) and after drug administration for flecainide 1.5 *µ*M (second row), vernakalant 10 *µ*M (third row) and tertiapin-Q 0.1 *µ*M (fourth row). In this case, vernakalant 10 *µ*M produced the greater drug-induced changes in DF and meandering area, followed by flecainide and, finally, tertiapin-Q. This illustrates how drugs tended to decelerate the rotor (reducing DF) and widen the rotor’s trajectory (increasing the meandering area).

**Fig 3.**
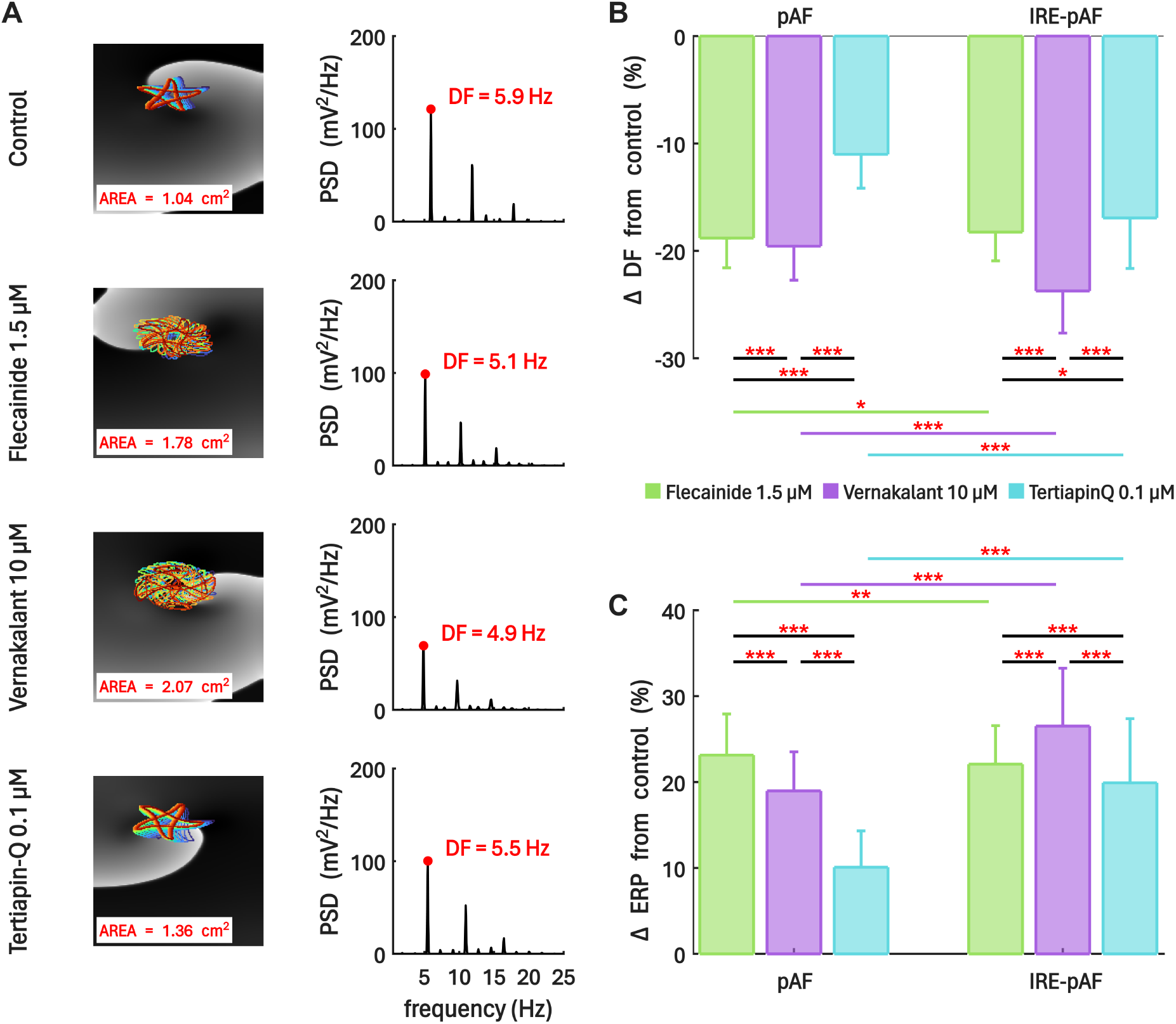
Drug-induced changes in rotor dominant frequency (DF) and ERP across the two pAF-like populations. A: Rotor tip trajectory and meandering area, and PSD and DF for a specific profile before drug application (top row) and after the application of flecainide 1.5 *µ*M (second row), vernakalant 10 *µ*M (third row), tertiapin-Q 0.1 *µ*M (fourth row), illustrating drug-induced rotor slowing and increased meandering area. B-C: Drug-induced changes in DF (B) and ERP (C) at low concentrations for both populations. Values are given as mean *±* SD. For DF analyses, only non-terminated rotors were considered, with values computed as the median per profile, whereas ERP analyses included all profiles. Statistical comparisons were performed using the Wilcoxon rank-sum test: \**p <* 0.05, \*\**p <* 0.01, \*\*\**p <* 0.001.

Electrophysiological Substrate-dependent differences in DF reduction were observed across drugs (Fig 3B). Vernakalant produced a larger DF reduction in IRE-pAF than in pAF (*−*23.7 *±* 3.9% vs *−*19.6 *±* 3.1%, *p <* 0.001) and a similar trend was observed for tertiapin-Q (*−*16.9 *±* 4.7% vs *−*11.0 *±* 3.1%, *p <* 0.001). In contrast, flecainide induced slightly lower DF reductions in IRE-pAF compared to the pAF population (*−*18.3 *±* 2.7% vs *−*18.8 *±* 2.8%, *p <* 0.05). Within each population, drugs exhibiting larger DF reductions also showed higher successful cardioversion rates. In pAF, vernakalant, flecainide and tertiapin-Q reduced DF by *−*19.6 *±* 3.1%, *−*18.8 *±* 2.8% and *−*11.0 *±* 3.1%, corresponding to termination rates of 41%, 34% and 11%, respectively (Fig 2B). A similar pattern was observed in the IRE-pAF population, where DF reductions of *−*23.7 *±* 3.9%, *−*18.3 *±* 2.7% and *−*18.3 *±* 2.7% were associated with cardioversion efficacies of 44%, 21% and 18% for vernakalant, flecainide and tertiapin-Q, respectively (Fig 2B). Thus, greater rotor deceleration was generally associated with higher cardioversion response within each electrophysiological substrate.

Drug-induced changes in rotor meandering area showed a similar electrophysiological substrate-dependent tendency (see S3 Figure). Vernakalant and tertiapin-Q produced a larger increase in rotor meandering area in the IRE-pAF population compared to pAF (+94.9 *±* 24.0% vs +82.4 *±* 27.1%, *p <* 0.001, for vernakalant; +80.7 *±* 25.6% vs +65.9 *±* 24.2%, *p <* 0.001, for tertiapin-Q), whereas flecainide exhibited no significant difference between substrates (+76.0 *±* 21.8% in pAF vs +77.1 *±* 22.0% in IRE-pAF, n.s.). Increased meandering area may contribute to rotor destabilization by promoting wider rotor trajectories and increasing the probability of collision with tissue boundaries, thereby facilitating termination. These findings suggest that, in addition to greater rotor deceleration, enhanced spatial destabilization of reentry may contribute to the improved performance of vernakalant and tertiapin-Q in the IRE-pAF substrate.

In contrast, ERP prolongation showed a partially distinct behavior (Fig 3C). In the IRE-pAF population, the trend of ERP prolongation followed the same qualitative order as cardioversion success rates. However, in pAF, flecainide induced a larger ERP increase than vernakalant (+23.1 *±* 4.8% vs +19.0 *±* 4.5%, *p <* 0.001) despite showing lower cardioversion efficacy (34% vs 41%). Importatly, electrophysiological substrate-dependent differences were also observed for ERP increase. While flecainide produced less ERP prolongation in the IRE-pAF population compared to pAF (+22.1 *±* 4.5% vs +23.1 *±* 4.8%, *p <* 0.01), vernakalant and tertiapin-Q had a greater effect increasing ERP in the IRE-pAF population compared to pAF (+26.5 *±* 6.7% vs +19.0 *±* 4.5%, *p <* 0.001, for vernakalant; +19.9 *±* 7.5% vs +10.1 *±* 4.2%, *p <* 0.01, for tertiapin-Q).

Drug-specific differences in DF slowing may partially arise from distinct effects on refractoriness and conduction properties. Although flecainide and vernakalant produced partially comparable ERP prolongation, vernakalant preserved CV more effectively (CV reduction of *−*5.9 *±* 0.8% in pAF and *−*5.6 *±* 0.7% in IRE-pAF) compared to flecainide (*−*9.6 *±* 0.9% in pAF and *−*9.5 *±* 0.8% in IRE-pAF), resulting in larger WL prolongation and, therefore, contributing to stronger rotor deceleration (drug-induced CV changes are shown in S4 Figure). In contrast, tertiapin-Q produced negligible effects on CV (+0.1 *±* 0.6% in pAF and +0.4 *±* 0.7% in IRE-pAF), suggesting that its WL prolongation arises predominantly from refractoriness changes rather than conduction effects. Supplementary analyses showed that drug-induced WL changes correlated consistently with rotor deceleration across drugs and substrates (see S5 Figure).

Similar qualitative trends in drug-induced changes in DF and ERP were observed at high concentration (see S2 Figure), supporting that the antiarrhythmic advantage of vernakalant and tertiapin-Q in IRE-pAF, may be more closely linked to their ability to decelerate reentry than to ERP prolongation alone.

### Rotor dominant frequency influences cardioversion outcome

We next examined whether rotor DF before drug administration influenced the likelihood of successful cardioversion.

For this purpose, models were stratified according to rotor DF into 0.5 Hz intervals. As shown in Fig 4 for high drug concentrations, cardioversion success progressively decreased as rotor DF increased across all simulated drugs and populations. However, this decline differed across drugs: vernakalant preserved higher efficacy than flecainide at elevated rotor frequencies and this separation became more pronounced in the IRE-pAF population. Accordingly, the efficacy difference between vernakalant and flecainide increased with DF, particularly under inward-rectifier enhancement (Fig 4B). In the highest frequency interval (6.5-7.3 Hz), flecainide efficacy decreased from 49% in pAF to 40% in IRE-pAF, whereas vernakalant efficacy increased from 56% to 76%. Consequently, the relative advantage of vernakalant over flecainide increased from 7 to 36 percentage points. Tertiapin-Q also showed greater efficacy in IRE-pAF within this interval (19% versus 4% in pAF), although its absolute efficacy remained lower than that of flecainide and vernakalant. These results show that increasing DF accentuated substrate-dependent differences in pharmacological response, with inward-rectifier enhancement favoring vernakalant and tertiapin-Q compared to flecainide. Similar trends were observed at low concentrations (S6 Figure), although overall cardioversion rates were lower.

**Fig 4.**
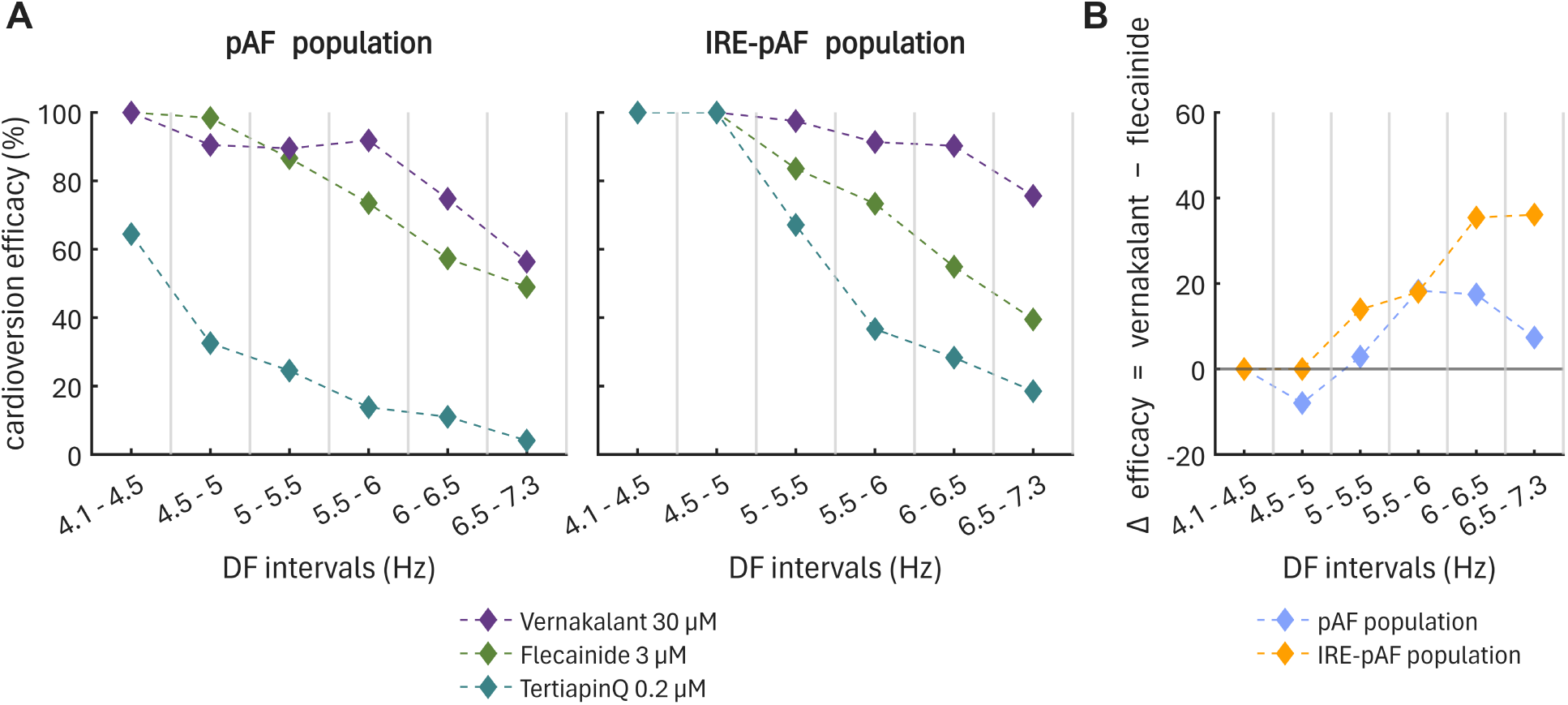
Cardioversion efficacy as a function of rotor dominant frequency (DF). A: Simulated cardioversion efficacy at high drug concentrations for the pAF and IRE-pAF populations. B: Difference in cardioversion efficacy between vernakalant and flecainide within each DF interval, calculated as vernakalant minus flecainide and expressed in percentage points (positive values indicate higher efficacy of vernakalant). DF is stratified into 0.5-Hz intervals.

### High-dominant frequency rotors reveal divergent drug efficacy and underlying mechanisms of cardioversion

To further explore the impact of rotor DF on drug response, sustained rotors were stratified into extreme low- and high-frequency rotor subgroups and we examined cardioversion efficacy and drug-induced changes, as shown in Fig 5 for high concentration (see S7 Figure for low concentration analysis).

**Fig 5.**
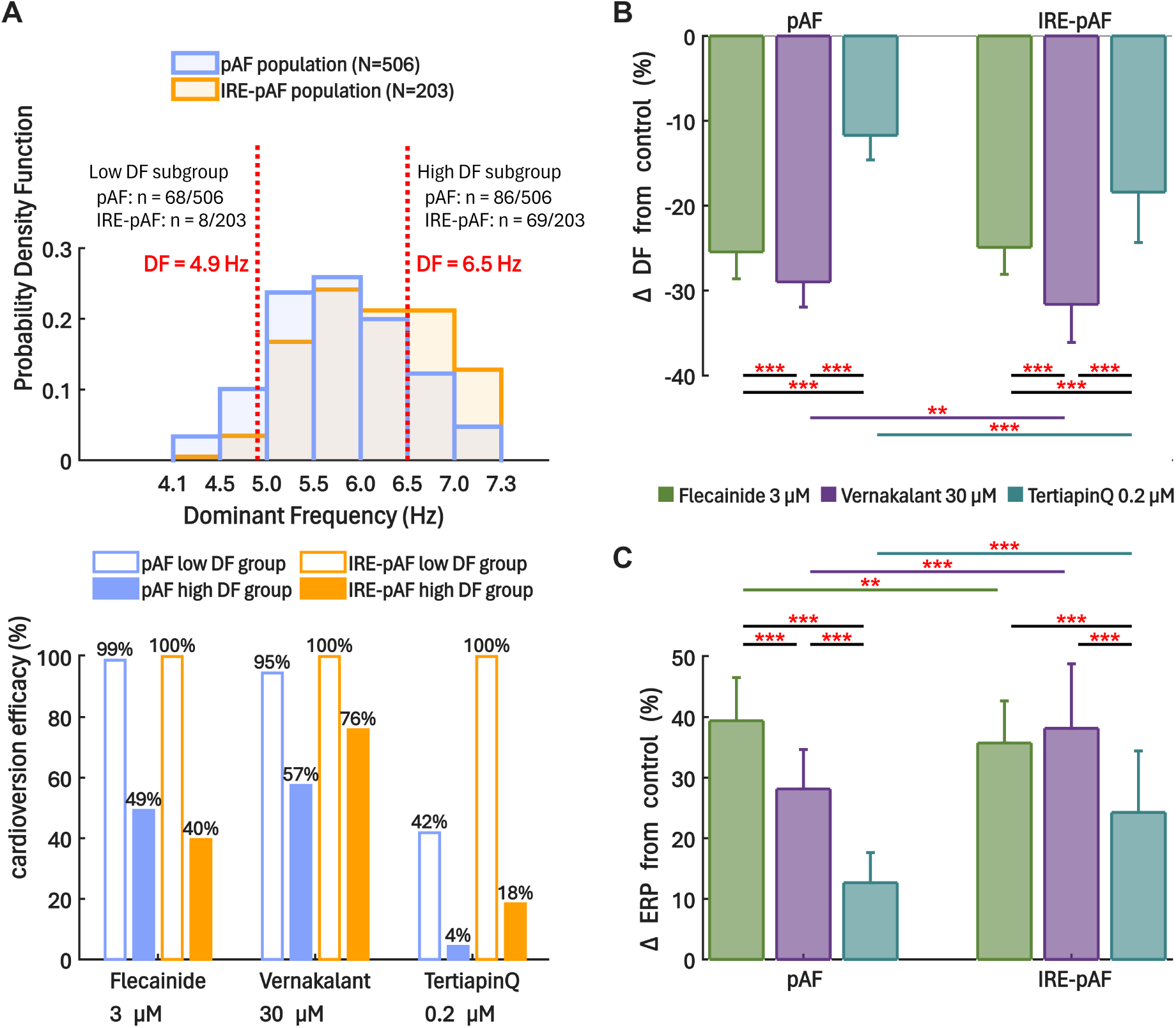
Cardioversion efficacy and electrophysiological responses in low- and high-frequency rotor subgroups. A: Top panel, distribution of rotor dominant frequency (DF) across the population, illustrating the stratification into low- and high-frequency subgroups using predefined thresholds (top panel); Bottom panel, cardioversion efficacy for each drug within the defined DF subgroups (blank and filled bars for low- and high-subgroups, respectively). Cardioversion was defined as termination of sustained rotor activity during the simulation following drug administration and computed as the ratio of terminated rotors to total sustained rotors within each subgroup. B-C: Drug-induced changes in DF (B) and ERP (C) at high concentrations for the high-frequency subgroup, shown for both pAF and IRE-pAF populations. Values are given as mean *±* SD. For DF analyses, only non-terminated rotors were considered, with values computed as the median per profile, whereas ERP analyses included all profiles. Statistical comparisons were performed using the Wilcoxon rank-sum test: \**p <* 0.05, \*\**p <* 0.01, \*\*\**p <* 0.001.

As represented in Fig 5A-top panel, ionic profiles were stratified into low- and high-DF subgroups using the mean *±* SD thresholds of the clinical DF distribution reported by Atienza et al. [11]. This resulted in a low DF subgroup (4.1–4.9 Hz; n = 68 in pAF and n = 8 in IRE-pAF) and a high DF subgroup (6.5–7.3 Hz; n = 86 in pAF and n = 69 in IRE-pAF). This stratification enabled the comparison of drug cardioversion success and electrophysiological response across distinct dynamical regimes. Importantly, high-DF models were significantly more frequent in the IRE-pAF population than in pAF (69/203, 34.0%, vs 86/506, 17.0%; Fisher’s exact test, *p <* 0.001), indicating that inward-rectifier enhancement increased the likelihood of high-frequency reentrant dynamics.

Cardioversion efficacy differed markedly across DF subgroups (Fig 5A-bottom pannel). In the low-frequency subgroup, all drugs achieved high cardioversion rates, with only modest differences between compounds and populations. In contrast, in the high-frequency subgroup, termination rates were substantially reduced and drug-specific differences became more pronounced. In pAF, flecainide 3*µ*M decreased from 99% cardioversion efficacy in the low-DF subgroup to 49% in the high-DF subgroup, whereas vernakalant 30*µ*M declined from 95% to 57%. In IRE-pAF, the deterioration was even more marked: flecainide dropped from 100% to 40%, whereas vernakalant decreased from 100% to 76%. Tertiapin-Q 0.2*µ*M also showed a strong dependence on DF, especially in IRE-pAF, where performance fell from 100% to 18%. These results indicated that high rotor frequency is a major determinant of reduced pharmacological success, but also that vernakalant preserves efficacy substantially better than flecainide under the fastest dynamical conditions. Similar trends were observed at low concentrations (see S7 Figure).

To further investigate the mechanisms underlying these differences, we analyzed drug-induced changes in DF and ERP within the high-frequency subgroup (Fig 5B-C). This analysis was restricted to high DF rotors, as this regime exhibited the greatest divergence in cardioversion efficacy and therefore provided the most informative setting to identify the mechanistic determinants of successful termination. Flecainide produced a comparable DF reduction in pAF and IRE-pAF (approximately *−*25% *±* 3% in both, n.s.), whereas vernakalant and tertiapin-Q decelerated rotors more strongly in IRE-pAF than in pAF (*−*31.6 *±* 4.5% vs *−*29.0 *±* 3.0%, *p <* 0.001, for vernakalant; *−*18.4 *±* 5.9% vs *−*11.7 *±* 2.9%, *p <* 0.001, for tertiapin-Q). Within each population, vernakalant induced the highest decrement in DF, followed by flecainide and tertiapin-Q, following the same trend as cardioversion success rates. By contrast, ERP changes did not reproduce the same trend across drugs and populations. Significant electrophysiological substrate-dependent differences in ERP prolongation were observed: flecainide showed a higher impact in the pAF population (+39.4 *±* 7.1% vs +35.7 *±* 6.9%, *p <* 0.01) while vernakalant and tertiapin-Q induced greater change in the IRE-pAF population (+38.1 *±* 10.6% vs +28.1 *±* 6.5%, *p <* 0.001, for vernakalant; +24.3 *±* 10.2% vs +12.7 *±* 5.0%, *p <* 0.001, for tertiapin-Q). Notably, flecainide caused the largest ERP prolongation in the pAF population, whereas vernakalant did so in the IRE-pAF population. However, in both populations, vernakalant exhibited greater termination rates. Tertiapin-Q induced the lowest ERP prolongation in both populations, matching its lowest efficacy in cardioversion. Hence, even among high-frequency rotors, the therapies that preserved cardioversion performance more effectively in IRE-pAF were those that reduced rotor DF more strongly.

### Distinct ionic signatures underlie pharmacological cardioversion and drug-specific response

Rotor dynamics were closely associated with cardioversion efficacy, but the ionic basis underlying these responses remains unclear. To address this, we analyzed the ionic signatures of responder models and drug-specific exclusive responders across drugs and populations. Fig 6 summarizes the results obtained at low drug concentrations, while the corresponding analyses at high concentrations is provided in S8 Figure.

**Fig 6.**
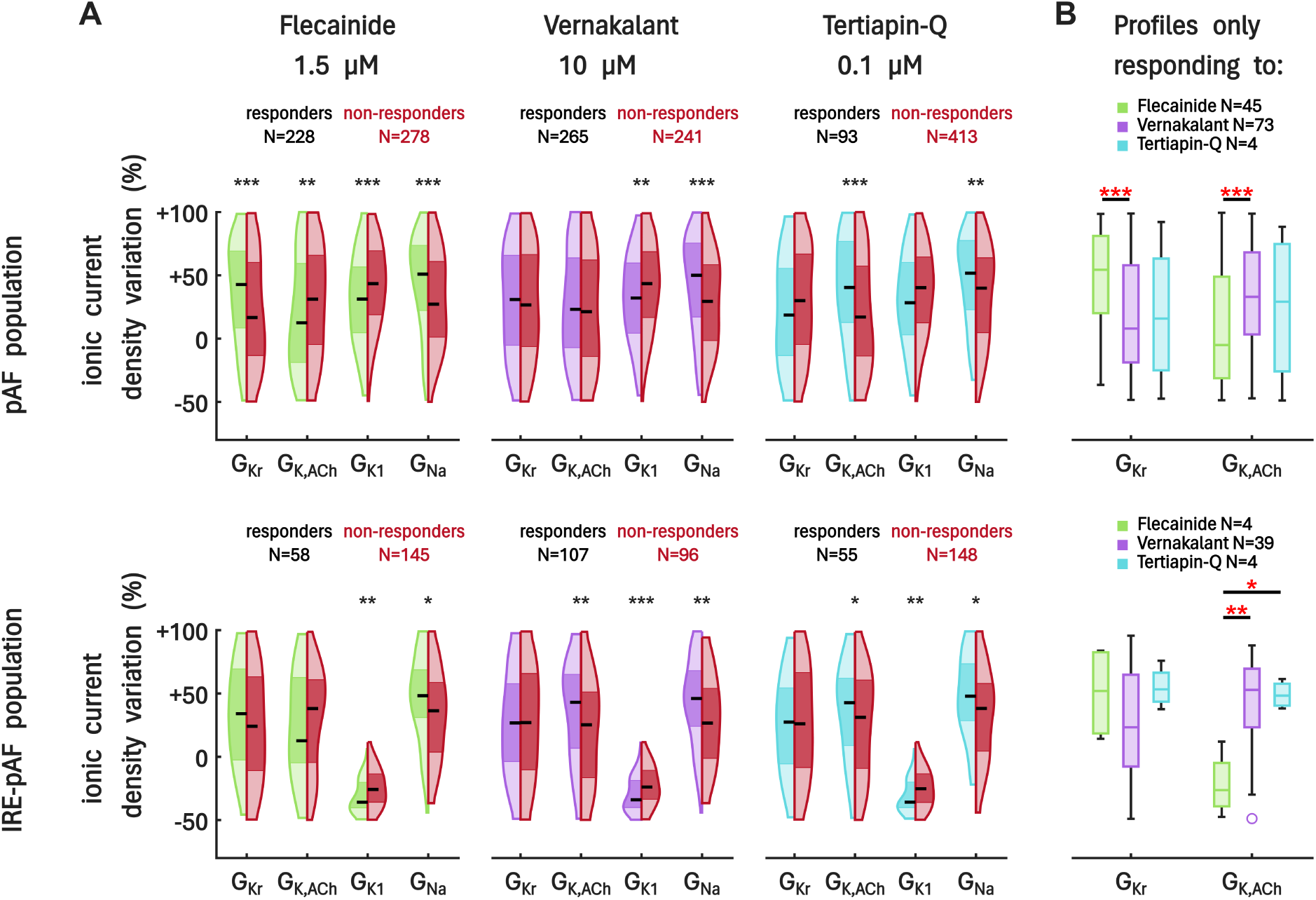
Ionic signatures of pharmacological cardioversion. A: Distribution of selected ionic conductances (G_Kr_, G_K,ACh_, G_K1_, and G_Na_) for responder and non-responder models across drugs and populations. Top panels correspond to the pAF population and bottom panels to the IRE-pAF population. Responders were defined as profiles in which pharmacological intervention terminated sustained rotor activity. B: Distribution of selected ionic conductances (G_Kr_, G_K,ACh_) for profiles exclusively cardioverted by each drug for pAF (top panel) and IRE-pAF (bottom panel) populations. Statistical comparisons were performed using the Wilcoxon rank-sum test: \**p <* 0.05, \*\**p <* 0.01, \*\*\**p <* 0.001.

Responder and non-responder ionic profiles showed distinct distributions across drugs and substrates (Fig 6A). In the pAF population (n = 506 profiles exhibiting sustained reentry), drug action induced a favorable response in 228 models for flecainide 1.5*µ*M (45%), 265 for vernakalant 10*µ*M (52%) and 93 for tertiapin-Q 0.1*µ*M (18%). In the IRE-pAF population (n = 203), responder models were 58 (29%) for flecainide 1.5*µ*M, 107 (53%) for vernakalant 10*µ*M and 55 (27%) for tertiapin-Q 0.1*µ*M. These results indicate that vernakalant maintains consistent efficacy across substrates, whereas flecainide shows a marked reduction in response in the IRE-pAF population. In contrast, tertiapin-Q increases its responder rate under inward-rectifier enhancement, which is consistent with the observed electrophysiological substrate-dependent improvement in cardioversion success.

These substrate- and drug-dependent differences were further investigated by evaluating the distinct ionic properties between responder and non-responder models. Across all drugs and populations, inward-rectifier current I_K1_ and sodium current I_Na_ exhibited the most consistent pattern with non-responders showing significantly higher G_K1_ and lower G_Na_ compared to responders.

By contrast, I_K,ACh_ and I_Kr_ exhibited drug-specific patterns. Responders to tertiapin-Q consistently showed higher G_K,ACh_ in both populations, in line with its selective blockade of this current. Vernakalant showed a similar dependence in the IRE-pAF population, where I_K,ACh_ is upregulated, whereas flecainide responders tended to exhibit lower G_K,ACh_, particularly in the pAF population. These opposing trends indicate that the role of I_K,ACh_ in cardioversion is strongly drug-dependent. Differences in I_Kr_ were primarily observed for flecainide, with responders exhibiting higher G_Kr_. In addition, other currents associated with repolarization reserve and membrane stability, such as I_to_, I_SK_ and I_K2P_, tended to be elevated in non-responders (see S9 Figure), reinforcing the association between enhanced repolarization capacity and resistance to cardioversion. Other repolarization currents, including I_Kur_ and I_Ks_, did not show consistent differences between responders and non-responders across drugs and populations (see S9 Figure), suggesting that these currents play a limited role in determining pharmacological response.

To further characterize drug-specific response, we analyzed models that were exclusively cardioverted by a single drug (Fig 6B). In the pAF population, 45 models responded exclusively to flecainide 1.5*µ*M, 73 to vernakalant 10*µ*M and 4 to tertiapin-Q 0.1*µ*M. In the IRE-pAF population, exclusive responders to flecainide decreased markedly to 4 models, whereas vernakalant retained 39 exclusive responders and tertiapin-Q remained limited (4 models). This shift suggests that inward-rectifier enhancement reduces the number of profiles uniquely responsive to flecainide, while preserving a larger subset of models preferentially responsive to vernakalant. Profiles exclusively responding to flecainide tended to exhibit higher G_Kr_ and lower G_K,ACh_, suggesting a potential dependence on repolarization pathways sensitive to I_Na_/I_Kr_ blockade. In contrast, profiles exclusively responding to vernakalant and tertiapin-Q showed higher G_K,ACh_, consistent with the role of I_K,ACh_ inhibition in mediating their antiarrhythmic effects. Notably, not all ionic differences aligned with the primary targets of each drug, indicating that cardioversion susceptibility arises from multivariate interactions between ionic currents rather than single-current determinants. For instance, G_Kur_ tended to be higher in flecainide exclusive responders compared to vernakalant exclusive responders, despite vernakalant exerting a stronger block of this current (data not shown). Although this difference did not reach statistical significance, it further illustrates that drug efficacy cannot be directly inferred from single-current targeting.

## Discussion

The present simulation study provides new insights into how substrate-specific electrophysiology modulates cardioversion success and the mechanisms underlying pharmacological cardioversion across distinct pAF-like conditions. By incorporating two different levels of inward rectifier levels, we constructed two electrophysiological substrates representing pAF and IRE-pAF populations, which showed differential baseline arrhythmogenic characteristics emerging from tissue-level simulations, including differences in reentry inducibility, rotor stability and DF. Importantly, both populations were independently calibrated against the same experimental pAF datasets, representing one of the first population-based studies to incorporate calibration against pAF-specific electrophysiological data. For both populations, three antiarrhythmic drugs were simulated at two therapeutic concentrations, resulting in in-silico efficacy trends broadly consistent with reported clinical cardioversion rates. The main findings are: (i) electrophysiological substrate fundamentally reshapes pharmacological response patterns, leading to differential efficacy across antiarrhythmic strategies; (ii) inward-rectifier enhancement penalizes flecainide efficacy while preserving or favoring I_K,ACh_-targeting strategies (vernakalant and tertiapin-Q); (iii) pharmacological cardioversion efficacy progressively declined with increasing rotor DF; (iv) inward-rectifier enhancement increased the prevalence of high-DF rotors, where vernakalant showed a progressively larger efficacy advantage over flecainide; (v) successful cardioversion was more closely associated with drug-induced rotor deceleration than with refractoriness prolongation alone. Together, these findings support that antiarrhythmic drug response emerges from the interaction among electrophysiological substrate, rotor dynamics and drug-specific mechanisms.

To our knowledge, this study represents one of the first population-based investigations specifically incorporating pharmacological I_K,ACh_-targeting mechanisms. While I_K,ACh_ modulation has been extensively investigated experimentally, its integration within antiarrhythmic simulations remains comparatively limited. Our findings suggest that incorporating pharmacological targeting of I_K,ACh_ contributes to understanding substrate-dependent drug interactions and pharmacological variability across AF substrates.

Finally, our work reinforces the value of population-based computational frameworks to investigate variability in antiarrhythmic drug response and highlights the potential of in-silico approaches to support the development of DF- and substrate-guided pharmacological strategies and future personalized approaches for AF management.

### AF electrophysiological substrate reshapes antiarrhythmic efficacy and defines pharmacological response patterns

Our results show that the underlying electrophysiological substrate had a marked impact on antiarrhythmic efficacy, leading to distinct pharmacological response patterns across the two pAF-like populations. Despite being calibrated to the same biomarkers, the pAF and IRE-pAF populations exhibited different arrhythmogenic characteristics, including differences in refractoriness, rotor DF and vulnerability to reentry.

Notably, while effective G_K1_ distributions largely overlapped between populations after calibration, effective G_K,ACh_ remained markedly shifted toward higher values in IRE-pAF. Therefore, although both inward rectifier currents were enhanced in the baseline formulation, the final calibrated populations differed more prominently in their effective I_K,ACh_ contribution, suggesting that part of the observed electrophysiological substrate-dependent effects may be driven by cholinergic inward rectifier activity. Consistent with this observation, Atienza et al. [11] demonstrated that activation of adenosine-modulated I_K,ACh_ shortens atrial refractoriness, accelerates DF and promotes stabilization of reentrant activity and AF maintenance. Thus, the greater contribution of I_K,ACh_ in the IRE-pAF population likely contributed to refractoriness shortening and DF acceleration.

These population-specific properties translated into differential cardioversion rates across drugs. Importantly, the overall efficacy hierarchy observed in our simulations remained broadly consistent with previous clinical and computational studies. Vernakalant generally outperformed flecainide in recent-onset AF clinical cohorts (67 vs 46% [55] and 86 vs 78% [56]), while previous population-based simulations in structurally healthy atria similarly reported superior performance of vernakalant (65%) compared with flecainide (56%) in terminating AF episodes [5].

Beyond reproducing overall successful cardioversion trends, a major observation of the present work was that the underlying electrophysiological substrate differentially modulated pharmacological response. Inward-rectifier enhancement reduced flecainide efficacy while preserving and, in some cases, improving the performance of vernakalant and tertiapin-Q despite the more arrhythmogenic characteristics of the IRE-pAF population. The deterioration observed for flecainide performance under inward-rectifier enhancement is in agreement with previous observations linking higher AF complexity and faster activation rates with reduced cardioversion success [19, 20]. Thus, the faster and more stable rotor dynamics emerging under inward-rectifier enhancement may contribute to the reduced efficacy of conventional I_Na_/I_Kr_-mediated compounds such as flecainide. In contrast, drugs involving I_K,ACh_ targeting (vernakalant and tertiapin-Q) showed relative preservation under the more arrhythmogenic conditions of the IRE-pAF population. I_K,ACh_ has emerged as an attractive atrial-selective therapeutic target because of its role in reducing AF burden [57, 58] and its limited ventricular contribution [10]. Consistent with these observations, the selective blockade of I_K,ACh_ by tertiapin-Q was sufficient to counteract the expected response deterioration associated with inward-rectifier enhancement, while vernakalant preserved and even slightly improved cardioversion efficacy under the same conditions. Interestingly, previous studies investigating the selective inhibition of I_K,ACh_ have reported mixed results. Experimental studies demonstrated that tertiapin-Q reduced AF inducibility in neonatal rat whole-heart preparations [57], while the selective I_K,ACh_ inhibitor XAF-1407 showed promising antiarrhythmic efficacy in preclinical models [58]. In contrast, the first clinical evaluation of an I_K,ACh_-targeting strategy failed to demonstrate a reduction in AF burden compared with placebo in patients with pAF [59]. Our results may help reconcile these observations by suggesting that the antiarrhythmic performance of I_K,ACh_-targeting therapies cannot be considered independently of the underlying electrophysiological substrate.

Consistent with these observations, the ionic analysis further supported the hypothesis that substrates exhibiting enhanced I_K,ACh_ activity may be particularly susceptible to pharmacological strategies involving I_K,ACh_ targeting. Responders to vernakalant and tertiapin-Q consistently exhibited higher G_K,ACh_ values than non-responders, while profiles responding exclusively to these therapies also exhibited up-regulated G_K,ACh_. In contrast, flecainide-exclusive responders tended to exhibit lower G_K,ACh_ together with higher G_Kr_, consistent with previous observations linking repolarization reserve to flecainide efficacy [5]. The enrichment of G_K,ACh_ observed both at the population level and among vernakalant and tertiapin-Q responsive profiles highlights the contribution of cholinergic inward rectifier activity to electrophysiological substrate-specific pharmacological susceptibility.

However, selective I_K,ACh_ inhibition alone appeared insufficient to reproduce the efficacy achieved by broader atrial-selective multichannel targeting. Despite producing comparable high-concentration I_K,ACh_ blockade, tertiapin-Q consistently remained less effective than vernakalant across electrophysiological substrates. Previous computational studies investigating vernakalant pharmacology suggested that its antiarrhythmic action emerges from the integrated modulation of multiple ionic mechanisms rather than isolated channel blockade [30, 31]. More recently, Dasi et al. [60] demonstrated that simultaneous targeting of multiple atrial-selective ionic currents may produce synergistic antiarrhythmic effects beyond those expected from individual channel inhibition alone. Our findings further support this concept, indicating that while I_K,ACh_ blockade contributes substantially to electrophysiological substrate-specific efficacy preservation, broader atrial-selective multichannel targeting appears to remain important for achieving optimal cardioversion success.

### Rotor dominant frequency dynamics provide mechanistic insight into pharmacological cardioversion across drugs and substrates

Our study shows that rotor DF strongly influenced pharmacological responses across drugs and substrates. Higher DF values were consistently associated with lower cardioversion efficacy, whereas slower rotors were substantially more susceptible to pharmacological termination. These observations are in line with previous experimental and clinical studies linking lower atrial activation rates to favorable cardioversion outcomes [18], while elevated AF frequencies have been associated with increased AF complexity, progressive electrical remodeling and resistance to rhythm control interventions [8, 16–18]. Similarly, Bollmann et al. demonstrated that lower atrial fibrillatory rates (*≈* 5.6 Hz) predicted successful cardioversion with classic I_Na_/I_Kr_ blockers following oral flecainide [20] and intravenous ibutilide [19]. In our simulations, differences between pharmacological strategies became increasingly pronounced at higher frequencies, where vernakalant preserved efficacy more effectively than flecainide. Consistently, Mochalina et al. reported no significant differences in atrial fibrillatory rate between responders and non-responders undergoing vernakalant treatment, suggesting that vernakalant may remain effective across a broader range of activation frequencies [61]. Interestingly, inward-rectifier enhancement significantly increased the prevalence of high-DF reentrant dynamics, where the relative efficacy advantage of vernakalant over flecainide became progressively larger. Since rotor DF can be estimated clinically, these findings suggest that DF may also identify electrophysiological substrates with distinct expected responses to antiarrhythmic drugs. Rather than acting solely as a predictor of cardioversion success, DF may therefore represent a clinically accessible biomarker of substrate-dependent pharmacological susceptibility that could contribute to DF-guided antiarrhythmic drug selection.

Beyond rotor DF, drug-induced rotor deceleration provided a more consistent mechanistic correlate of cardioversion than refractoriness prolongation alone. Both ERP prolongation and deceleration of activation frequency have long been recognized as important antiarrhythmic mechanisms contributing to AF termination [45, 62, 63]. However, while all drugs increased refractoriness, ERP changes did not systematically follow cardioversion efficacy. For example, flecainide produced a larger ERP increase than vernakalant in the pAF population (+23.1 *±* 4.8% vs +19.0 *±* 4.5%) despite exhibiting lower cardioversion rates (34% vs 41%). In contrast, drug-induced DF reduction closely paralleled cardioversion outcomes across drugs and substrates. These observations are consistent with experimental evidence demonstrating that the magnitude of DF reduction is closely associated with pharmacological AF termination [63].

The relationship between rotor deceleration and cardioversion may be partially explained by the interaction between refractoriness and conduction properties. Activation frequency is inversely related to wavelength during reentrant activity, which itself depends on both refractoriness and conduction velocity. Consequently, antiarrhythmic interventions that increase wavelength tend to slow reentrant activity, enlarge rotor core dimensions and facilitate rotor destabilization and termination [62]. In our simulations, although flecainide and vernakalant both prolonged ERP, vernakalant preserved conduction velocity more effectively, resulting in greater wavelength prolongation and stronger rotor deceleration despite a smaller ERP increase. Previous experimental and computational studies similarly reported that both flecainide and vernakalant prolong atrial refractoriness, although vernakalant produces substantially less conduction slowing than flecainide [5, 23, 31, 64, 65]. This combination results in greater wavelength prolongation and consequently greater deceleration of reentrant activity. Consistent with this concept, van Hunnik et al. [66] demonstrated that vernakalant produced greater AF cycle length prolongation than flecainide in electrically remodeled goat atria, supporting greater deceleration of AF activation dynamics. Additionally, previous investigations have shown that flecainide markedly depresses conduction velocity compared with alternative antiarrhythmic strategies, including I_K1_ blockade with chloroquine [67] and I_Kr_ blockade [68]. Although effective in prolonging refractoriness, excessive conduction slowing may partially offset wavelength prolongation, limiting rotor deceleration and reducing cardioversion efficacy. These experimental observations are consistent with our simulations, where vernakalant achieved stronger DF reduction (i.e. greater wavelength prolongation) despite producing smaller ERP increases than flecainide.

The same mechanistic framework may also explain the behavior of selective I_K,ACh_ blockade. Experimental studies demonstrated that tertiapin-Q prolongs atrial refractoriness, reduces rotor DF and destabilizes reentrant activity [57]. In our simulations, tertiapin-Q produced larger ERP increases and stronger DF reduction in the IRE-pAF population, suggesting that selective I_K,ACh_ blockade may promote AF termination through rotor deceleration and destabilization.

Although DF emerged as the electrophysiological variable most consistently associated with cardioversion efficacy, additional markers of rotor stability may provide complementary insights into the mechanisms underlying rotor termination. Rotor destabilization likely involves mechanisms beyond DF deceleration alone. In addition to reducing DF, all three drugs increased rotor meandering area, while vernakalant and tertiapin-Q exhibited a particularly pronounced effect in the IRE-pAF population. Greater rotor meandering has been associated with reduced rotor stability and enhanced interaction with surrounding tissue boundaries [45, 57, 62, 67], potentially facilitating termination of reentrant activity.

Therefore, these findings suggest that pharmacological cardioversion is primarily associated with rotor DF reduction, whereas changes in the spatial organization of reentry may further facilitate AF termination.

### Limitations and future perspectives

Simulations presented here were performed in a simplified two-dimensional homogeneous tissue, which does not fully reproduce the anatomical complexity of the human atria, an important determinant of AF maintenance. Although this approach enabled systematic investigation of rotor dynamics and pharmacological response mechanisms, additional structural features such as patient-specific anatomy, fiber orientation and regional electrophysiological heterogeneity may further influence AF maintenance and antiarrhythmic efficacy. Nevertheless, previous three-dimensional biatrial computational studies have suggested that the ionic substrate may exert a greater influence on antiarrhythmic drug response than the extent or spatial distribution of structural remodeling [28]. Future studies should therefore extend the present framework toward anatomically realistic three-dimensional atrial models to evaluate whether the mechanistic relationships identified here remain consistent under more complex propagation conditions.

Additionally, pharmacological effects were represented using simple pore-block formulations. While this approach reproduces the principal ionic targets of each drug, it does not capture state-dependent channel kinetics that may influence excitability, conduction velocity and wavelength dynamics. In particular, previous studies have developed Markov formulations describing I_Na_ interactions with flecainide and vernakalant [31]. Incorporating such formalism may provide a more detailed characterization of drug-induced conduction changes and further refine mechanistic insights regarding rotor deceleration and cardioversion.

Finally, validation was performed through comparison with previously reported experimental and clinical observations rather than direct patient-specific data. Future integration of personalized computational models with clinical ECG or ECGi recordings may enable prospective evaluation of the substrate-dependent mechanisms identified here and facilitate the development of individualized rhythm-control strategies. In this context, the differential efficacy observed across substrates and drugs suggests that computational approaches may ultimately contribute to DF- and substrate-guided pharmacological selection. Our findings suggest that DF may represent a potentially useful clinical marker of underlying electrophysiological substrate properties and pharmacological susceptibility, potentially helping identify patients more likely to benefit from specific antiarrhythmic strategies.

## Conclusion

In pAF-like atrial models, antiarrhythmic efficacy depended on the interaction between electrophysiological substrate, rotor dynamics and drug mechanism. Inward-rectifier enhancement impaired flecainide response while preserving or improving I_K,ACh_-targeting strategies and shifted the response toward vernakalant at high baseline DF. These findings support DF as a marker of pharmacological susceptibility and motivate substrate-informed antiarrhythmic drug selection.

## Supporting information

**S1 Appendix. Supplementary methods.** Extended methodological details including baseline model description, calibration specifications and drug model generation.

**S1 Data. Calibrated pAF population.** Scaling factors for the 13 ionic conductances defining the 853 calibrated profiles included in the pAF population.

**S2 Data. Calibrated IRE-pAF population.** Scaling factors for the 13 ionic conductances defining the 282 calibrated profiles included in the IRE-pAF population.

**S1 Figure. Ionic variability across calibrated populations.** Distribution of effective conductance scaling factors for all variable ionic currents in the calibrated pAF (blue, N = 853) and IRE-pAF (orange, N = 282) populations. Red dashed lines indicate the parameter range limits used during population generation.

**S2 Figure. Drug-induced changes in rotor DF and ERP at high concentration.** Percentage variation (mean *±* SD) in rotor DF and ERP at high concentration for both populations. Statistical comparisons were performed using the Wilcoxon rank-sum test: *p*<*0.05, **p*<*0.01, ***p*<*0.001.

**S3 Figure. Drug-induced changes in the meandering area.** Percentage variation (mean *±* SD) in meandering area at low and high concentrations for both populations. Statistical comparisons were performed using the Wilcoxon rank-sum test: *p*<*0.05, **p*<*0.01, ***p*<*0.001.

**S4 Figure. Drug-induced changes in conduction velocity.** Percentage variation (mean *±* SD) in CV at low and high concentrations for both populations. Statistical comparisons were performed using the Wilcoxon rank-sum test: *p*<*0.05, **p*<*0.01, ***p*<*0.001.

**S5 Figure. Correlation analysis of drug-induced DF reduction and WL increase across drugs and substrates.** Spearman correlation between percentage changes in DF and WL following drug administration for flecainide, vernakalant and tertiapin-Q in both pAF and IRE-pAF populations. Each point corresponds to one ionic profile. Correlation coefficients (*ρ*) and p-values are indicated in each panel. Greater WL increases were generally associated with larger reductions in rotor frequency.

**S6 Figure. Cardioversion efficacy as a function of rotor dominant frequency for low concentration dosage.** A: Simulated cardioversion efficacy as a function of rotor dominant frequency (DF), stratified into frequency intervals for the two pAF-like populations. B: Difference in cardioversion efficacy between vernakalant and flecainide within each DF interval.

**S7 Figure. Cardioversion efficacy and electrophysiological responses in low- and high-frequency rotor subgroups at low concentration.** Cardioversion efficacy for each drug within the defined DF subgroups (A) and drug-induced changes in ERP (mean ± SD) and rotor DF (mean ± SD) for the high-frequency subgroup, shown for both pAF and IRE-pAF populations. Statistical comparisons were performed using the Wilcoxon rank-sum test: *p*<*0.05, **p*<*0.01, ***p*<*0.001.

**S8 Figure. Ionic signatures of pharmacological cardioversion under high concentration.** Distribution of selected ionic conductances (G_Kr_, G_K,ACh_, G_K1_, and G_Na_) for responder and non-responder models across drugs and populations (A) and distribution of selected ionic conductances (G_Kr_, G_K,ACh_) for profiles exclusively cardioverted by each drug (B). Statistical comparisons were performed using the Wilcoxon rank-sum test: *p*<*0.05, **p*<*0.01, ***p*<*0.001.

**S9 Figure. Comparison of responder and non-responder models across drugs and populations for repolarization currents.** Distribution of ionic conductance variation for G_SK_, G_K2P_, G_to_, G_Kur_ and G_Ks_. Statistical comparisons were performed using the Wilcoxon rank-sum test: *p*<*0.05, **p*<*0.01, ***p*<*0.001.

## Acknowledgments

This work was partially supported by the Dirección General de Política Científica de la Generalitat Valenciana (CIPROM/2023/14) and by grant PID2022-140553OB-C41 funded by MICIU/AEI/10.13039/501100011033 and by ERDF/EU. This project has also received funding from the European Union’s Horizon 2020 research and innovation program under grant agreement No 101016496 (SimCardioTest). The work in DF-R laboratory was partially supported by the Ministry of Science and Innovation (MCIN) (PID2023-150456OB-I00) funded by MCIN / AEI / 10.13039/501100011033). The Centro Nacional de Investigaciones Cardiovasculares (CNIC) is supported by the ISCIII, MCIN and the Pro CNIC Foundation, and is a Severo Ochoa Center of Excellence (CEX2020-001041-S funded by MICIN/AEI/10.13039/501100011033). The author thankfully acknowledges RES resources provided by Universitat de València in Tirant to IM-2026-1-0024 and Barcelona Supercomputing Center in MareNostrum5 to IM-2024-1-0010, IM-2024-2-0015, IM-2024-3-0001, IM-2025-1-0010, IM-2025-2-0007, IM-2025-3-0004 and IM-2026-1-0088.

